# SCODE: An efficient regulatory network inference algorithm from single-cell RNA-Seq during differentiation

**DOI:** 10.1101/088856

**Authors:** Hirotaka Matsumoto, Hisanori Kiryu, Chikara Furusawa, Minoru S.H. Ko, Shigeru B.H. Ko, Norio Gouda, Tetsutaro Hayash, Itoshi Nikaido

**Affiliations:** Bioinformatics Research Unit, Advanced Center for Computing and Communication, RIKEN, Saitama, Japan; Department of Computational Biology and Medical Sciences, Faculty of Frontier Sciences, The University of Tokyo, Chiba, Japan; Quantitative Biology Center (QBiC), RIKEN, Osaka, Japan; Universal Biology Institute, The University of Tokyo, Tokyo, Japan; Department of Systems Medicine, Keio University School of Medicine, Tokyo, Japan

## Abstract

The analysis of RNA-Seq data from individual differentiating cells enables us to reconstruct the differentiation process and the degree of differentiation (in pseudo-time) of each cell. Such analyses can reveal detailed expression dynamics and functional relationships for differentiation. To further elucidate differentiation processes, more insight into gene regulatory networks is required. The pseudo-time can be regarded as time information and, therefore, single-cell RNA-Seq data are time-course data with high time resolution. Although time-course data are useful for inferring networks, conventional inference algorithms for such data suffer from high time complexity when the number of samples and genes is large. Therefore, a novel algorithm is necessary to infer networks from single-cell RNA-Seq during differentiation.

In this study, we developed the novel and efficient algorithm SCODE to infer regulatory networks, based on ordinary differential equations. We applied SCODE to three single-cell RNA-Seq datasets and confirmed that SCODE can reconstruct observed expression dynamics. We evaluated SCODE by comparing its inferred networks with use of a DNaseI-footprint based network. The performance of SCODE was best for two of the datasets and nearly best for the remaining dataset. We also compared the runtimes and showed that the runtimes for SCODE are significantly shorter than for alternatives. Thus, our algorithm provides a promising approach for further single-cell differentiation analyses.

The R source code of SCODE is available at https://github.com/hmatsu1226/SCODE.

## 1. Introduction

Conventional bulk RNA-Seq reveals the average gene expression of an ensemble of cells, and therefore does not permit the analysis of detailed states of individual cells. With the advancement of single-cell RNA-Seq (scRNA-Seq), we can now quantify the expression of individual cells and analyze detailed differences among cells [1]. This enables several analyses such as the identification of cell types [2, 3], especially of rare cells [4, 5], and the estimation of cellular lineages [6, 7].

In analyses by scRNA-Seq, the reconstruction of cellular differentiation processes attracts attention as a novel approach to revealing differentiation mechanisms [8]. The differentiation process can be reconstructed using dimension reduction [9, 10] and stochastic processes [11], for example, and the degree of differentiation (in pseudo-time) of each cell is characterized by the position in the reconstructed process. By investigating the expression pattern in pseudo-time, genes can be clustered into multiple groups with different biological functions [9]. Moreover, the regulatory cascade of cellular state transitions, such as differentiation, can be inferred by comparing the timings of up‐ and down-regulation [11, 12, 13].

In addition, scRNA-Seq also enables the calculation of accurate correlations of expression between genes because scRNA-Seq can distinguish the detailed states of individual cells without contamination from multiple cell types. The accurate co-expression pattern of each cell type (progenitor cells and multiple types of differentiated cells) can reveal the key regulatory factors for lineage programming [14].

In this way, expression dynamics in pseudo-time and accurate relationships among genes can be inferred from scRNA-Seq data. For the next step in differentiation analyses using scRNA-Seq, it is important to reveal the regulatory interactions among genes that bring about the observed expression dynamics during differentiation, namely, gene regulatory network (GRN) inference from scRNA-Seq data. Pseudo-time can be regarded as time information, and hence, scRNA-Seq performed on cells undergoing differentiation can be regarded as time-course expression data at a high temporal resolution. Although several algorithms have been proposed to reconstruct GRN from time-course data [15], most of them are not suitable for scRNA-Seq data, such as that collected over continuous time and with a large number of samples. Moreover, time complexity is a serious problem, and runtime becomes infeasibly long with large numbers of samples and genes for the network inference from time-course data.

Recently, Boolean network-based algorithms have been proposed for inferring GRN from single-cell data [16, 17, 18]. Although these algorithms have revealed some interesting regulatory relationships, their time complexity increases significantly as the number of genes and cells increases, and they have thus been applied to data with a small number of genes. In addition, the expression data must first be converted into binary data for Boolean network inference, and therefore the relationship between networks and the underlying dynamics becomes obscured [19].

As another approach, ordinary differentiation equations (ODEs) have been used to describe regulatory network and expression dynamics. ODEs can describe continuous variables over continuous time and the underlying physical phenomena, and therefore they are suitable for inferring GRN from scRNA-Seq during differentiation. Although several ODE-based network-inference algorithms have been proposed [20, 21], most of them are not suitable for the differentiation case because these algorithms assume a steady-state condition. There are some ODE-based algorithms that infer GRNs such that the observed expression dynamics can be reconstructed from the optimized ODE [22]. However, time complexity is still a serious problem for such ODE-based algorithms [15]. Previous research has described optimizing an ODE by using single-cell data and pseudo-time to infer key GRNs [23]. Although it is a suggestive approach, the optimization assumes that the GRNs are given and learns the ODE for a specific GRN. Therefore, a novel and efficient algorithm is necessary to learn GRNs from ODEs designed for scRNA-Seq performed on differentiating cells and for a large number of samples and genes.

Accordingly, we developed an approach to describe regulatory networks and expression dynamics with linear ODEs as well as a novel, highly efficient optimization algorithm, SCODE, for scRNA-Seq performed on differentiating cells by integrating the transformation of linear ODEs and linear regression. In the Methods section, we show that linear ODEs can be transformed from fixed-parameter linear ODEs if they satisfy a relational expression. We also show that the relational expression can be estimated analytically and efficiently by linear regression. In addition, SCODE uses a small number of factors to reconstruct expression dynamics, which results in a marked reduction of time complexity. In the Results sections, we described the application of SCODE for three scRNA-Seq datasets during differentiation. First, we validated that the optimized ODEs can reconstruct observed expression dynamics accurately. Second, we evaluated the inferred network by comparing it to the transcription factor (TF) regulatory network database based on DNaseI footprints and transcription factor binding motifs. SCODE performed best with two of the datasets and was the close second best algorithm for the remaining dataset. Third, we compared the runtimes of the algorithms, and SCODE was significantly faster than previous algorithm that was designed for time-course data. Moreover, SCODE is faster than some algorithms that do not use time parameters. These results illustrate the remarkable efficiency of SCODE. Lastly, we analyzed the network inferred from a dataset and determined that the de novo methyltransferases *Dnmt3a* and *Dnmt3b* might be key regulators of differentiation.

In this paper, we propose a novel algorithm for scRNA-Seq performed on differentiating cells to reconstruct expression dynamics and infer regulatory networks with a highly efficient optimization method. We believe that our approach will substantially advance the development of regulatory network inference and promote the development of further single-cell differentiation analyses and bioinformatics methods.

## 2. Methods

### 2.1. Describing regulatory networks and expression dynamics with linear ODEs

In this research, we focus on TFs and inferring TF regulatory networks. First, we describe TF expression dynamics throughout differentiation with linear ODEs:

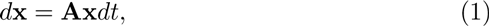

where **x** is a vector of length *G* (*G* is the number of TFs) that denotes the expression of TFs and **A** is a square matrix with dimensions equal to *G* that denotes the regulatory network among TFs. We infer the TF regulatory network by optimizing **A** such that the ODE can successfully describe the observed expression data.

The observed expression data consist of a *G* × *C* matrix (**X**^(*e*)^), where *C* is the number of cells. In addition, the time parameter of a cell *c* is given as *t_c_*. Therefore, our objective is to optimize **A** such that *d***x = Ax***dt* can properly represent **X**^(*e*)^ at a corresponding time point.

Here, **A** contains *G* × *G* parameters and an efficient parameter optimization algorithm is necessary for large values of *G*. This is because the time complexity is typically *O*(*G*^3^) for operation on a *G* × *G* matrix, and it will exceed *O*(*CG*^3^) to optimize **A** with a general algorithm. As experimental technologies have advanced, the number of cells that may be subjected to scRNA-Seq has been increasing, and hence *C* can be quite large. Therefore, we developed a novel algorithm to optimize **A** efficiently, even if both *G* and *C* are large, by integrating the transformation of linear ODEs and linear regression.

#### 2.1.1. Deriving **A** from a linear ODE transformation

At first, we consider the following linear ODE:

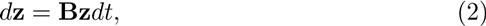

where **z** is a vector of length *G* and **B** is a known square matrix. If we know a matrix **W** that satisfies **x = Wz**, we can derive the ODE of **x** by transforming the ODE of **z** as follows:

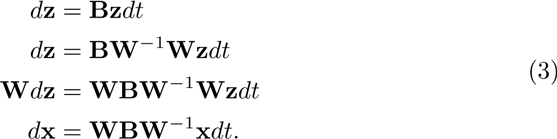

Therefore, if the parameter **B** of *d***z = Bz***dt* and the relationship **x = Wz** are given, we can derive **A** from **WBW**^−1^.

#### 2.1.2. Estimating **W** using linear regression

To infer **A**, we have to estimate a matrix **W** that satisfies **x = Wz**. Here, we assume that the problem of **W** inference can be regarded as a linear regression problem. Initially, from *d***z = Bz***dt*, we calculate **z** at *t = t_c_* for each cell and generate a *G* × *C* matrix (**Z**^(*e*)^) (Fig 1(a)). With this **Z**^(*e*)^, we optimize **W** to successfully represent the relationship **X**^(*e*)^ ≃ **WZ**^(*e*)^, which results in **x** ≃ **Wz**. The above problem can be regarded as solving the linear regression for each gene, as follows:

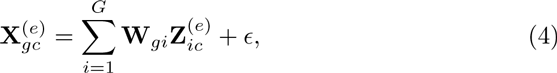

where *ϵ* is a noise term. Therefore, **W** can be optimized analytically and efficiently by linear regression for each TF (Fig 1(b)), and **A** can be efficiently calculated from **WBW**^−1^.

**Figure 1:**
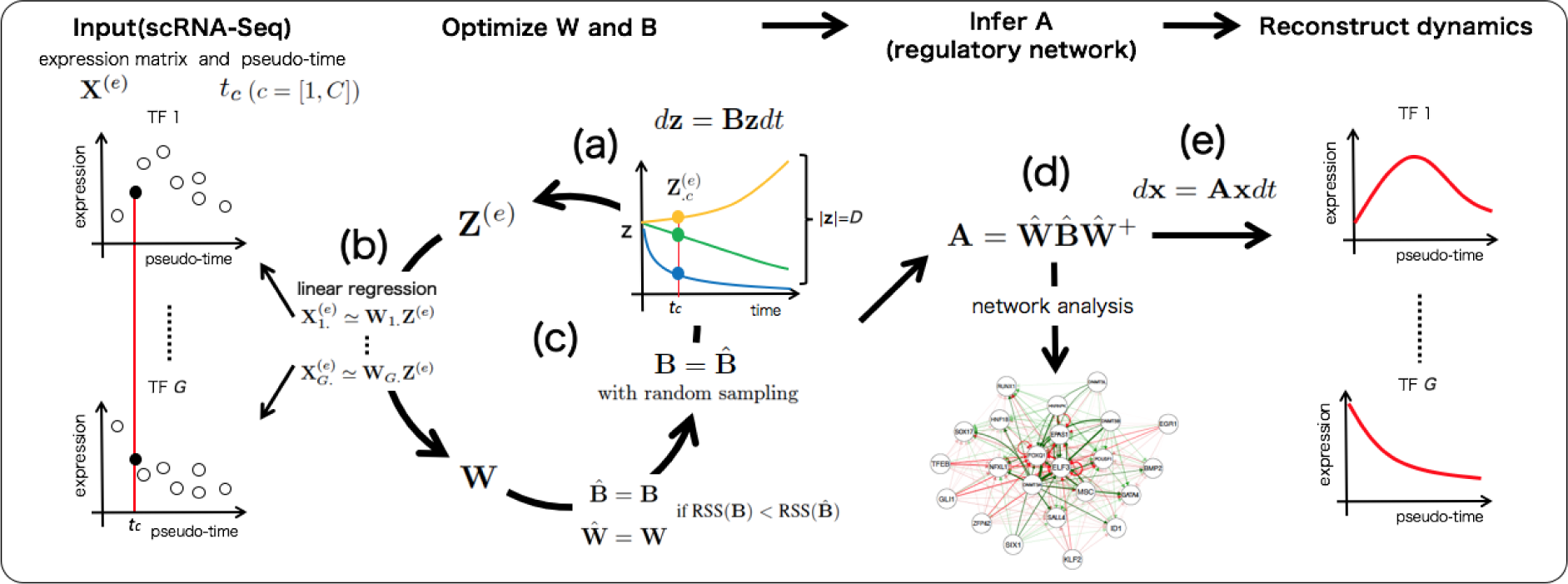
Abstract illustration of SCODE. (a) Sample **Z**^(*e*)^ from the ODE of **z** (b) Estimate **W** based on linear regression. (c) Optimize **B** iteratively. (d) Infer **A** from optimized **W** and **B**. (e) The expression dynamics can be reconstructed from the optimized ODE of **x**.

#### 2.1.3. Dimension reduction of **z**

The basic idea of reduction is that the patterns of expression dynamics are limited and expression dynamics can be reconstructed with a small number of patterns. For the next step, we consider a small vector **z** to represent the original expression dynamics. Hereafter, **z** is a vector of length *D*, with *D ≪ G*. In this case, **W** is a *G* × *D* matrix, and hence we used a pseudo-inverse matrix **W**^+^ instead of the inverse matrix, and **A** is derived from **A = WBW**^+^. The matrix **W** is estimated as before, via linear regression. By using a small vector **z**, the time complexity of estimation of **W** becomes much lower.

Recently, such dimensionality reduction approach has also been proposed to infer network [24]. Although it is a sophisticated algorithm, it is designed for discrete time-course data and small samples, and is not suitable for scRNA-Seq data.

#### 2.1.4. Optimizing **B**

Thus far, we have assumed **B** is given. To represent the original expression dynamics with small values of *D*, we optimize **B** for the next step. We suppose that the appropriate value of **B** satisfies the condition that the **Z**^(*e*)^ generated from *d***z = Bz***dt* can predict **X**^(*e*)^ with **WZ**^(*e*)^ accurately. Therefore, we evaluate the appropriateness of the matrix **B** with the following residual sum of squares (RSS):

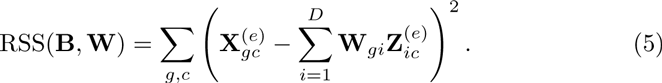

In this research, we assume **B** is a diagonal matrix and the elements **B***_ii_* satisfy *b*_min_ ≤ **B***_ii_* ≤ *b*_max_ (we set *b*_min_ and *b*_max_ to −10 and 2, respectively). This limitation is acceptable because large and small values of **B***_ii_* represent a dynamics of sharp change and seem to be an inefficient basis for reconstructing the expression dynamics.

We optimize **B** by random sampling and iterative optimization so that the RSS decreases (Fig 1(c)). The brief pseudocode is given below (see the supplementary text for the detailed procedure).

##### Algorithm 1 Iterative optimization of **B**

~~~
Initialize a diagonal matrix **B**^(1)^ randomly
**for** *k* = 1 : *I* **do**
  **Z**^(*e*)^ ⇐ Generate from *d***z = B**^(*k*)^**z***dt*
  **W**^(*k*)^ ⇐ Solution of linear regression (**X**^(*e*)^ ≃ **WZ**^(*e*)^)
  **if** 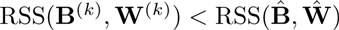 **then**
   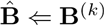
   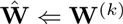
  **end if**
  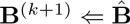
  *i* ⇐ uniform random value ∈ [1, *D*]
  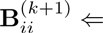 uniform random value ∈ [*b*_min_*, b*_max_]
**end for**
~~~

After the above optimization, **A** is inferred with 
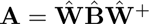
 (Fig 1(d)).

#### 2.1.5. Time complexity

The time complexity of optimizing **W** and **B** is 
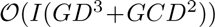
, where *I* is the number of iterations of **B** optimization. The time complexity of calculating **A** is 
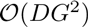
. Because we assume that *D* is small, the total time complexity is about 
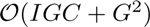
. As matrix operations on **A**, such as multiplication, have a time complexity of 
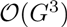
, our algorithm is highly efficient to infer regulatory network even though it integrates time-course information into the model.

### 2.2. Other network inference approaches

For comparison, we also developed a simple network inference algorithm based on linear regression that predicts expression of a particular TF from the expression of the remaining TFs as follows:

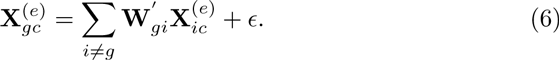

With this method, the optimized **W***′* is regarded as a regulatory network. In this research, we optimized **W***′* using two criteria. The first criterion is based on normal linear regression, and we estimated **W***′* with the *lm* function in R. The second criterion is based on lasso regression, and we estimated with the *msgps* package in R, which automatically selects the optimal degrees of freedom [25]. We used the Bayesian information criterion for model selection in *msgps*.

In addition, we inferred networks with GENIE3 [26], which also predicts TF expression from the expression of other TFs by using regression trees. The performance of GENIE3 was best in the DREAM5 Network Inference challenge for population data [27].

We also inferred networks with Jump3 [28], which is the expansion of GENIE3 for reconstructing a network from time-course expression data. Jump3 is based on jump trees and showed high performance for multiple time-course datasets.

### 2.3. Dataset

We analyzed three time-course scRNA-Seq datasets by the following procedures. First, transcripts per million reads (TPM) and fragments per millions of kilobases mapped (FPKM) were transformed as log(TPM+1) and log(FPKM+1), and we regarded these log-transformed values as the expression value. Next, we calculated the averaged expression of each TF at each time point and calculated the variance of the averaged expression for each TF. For TF data, we used Riken TFdb for mouse [29], and animalTFDB for human [30]. (Riken TFdb contains not only TFs but also their related genes, and we called all genes in the database transcription factors in this study.) Lastly, we regarded the TFs with large variances as variable TFs during differentiation. Hereafter, we used the top 100 variable TFs for network inference. For these 100 TFs, we estimated pseudo-time (*t_c_*) with Monocle [9]. We also excluded 100 randomly selected cells from the training data in order to use them as test data to evaluate adequate sizes of **z** (*D*).

#### 2.3.1. Data1: mouse ES cells to primitive endoderm cells

The first time-course scRNA-Seq dataset (at 0, 12, 24, 48, and 72 h) analyzed was derived from primitive endoderm (PrE) cells differentiated from mouse ES cells (by using G6GR ES cells [31]), containing 456 cells. This dataset was produced with RamDA-Seq, a novel scRNA-Seq protocol developed by our laboratory (in submission).

#### 2.3.2. Data2: mouse embryonic fibroblast cells to myocytes

The second dataset was derived from scRNA-Seq data obtained to examine direct reprogramming from mouse embryonic fibroblast (MEF) cells to myocytes at days 0, 2, 5, and 22 [32]. This dataset contained 405 cells.

#### 2.3.3. Data3: human ES cells to definitive endoderm cells

The third dataset was a scRNA-Seq time course (at 0, 12, 24, 36, 72, and 96 h) derived from definitive endoderm (DE) cells differentiated from human ES cells, containing 758 cells [33].

### 2.4. Network validation method

To validate the inferred networks, we used the Transcription Factor Regulatory Network database (http://www.regulatorynetworks.org), which was constructed from DNaseI footprints and TF-binding motifs [34, 35]. We integrated the TF regulatory networks of all cells for human and mouse, and extracted 100 × 100 TF regulatory networks for each dataset. We regarded these TF regulatory networks as correct networks for each dataset and calculated the AUC values of the inferred networks. The AUC values were calculated by regarding the directed edges that show higher absolute values as representing reliable regulatory relationships. We removed self-loop regulation and TFs that do not have an edge in the correct network from AUC calculation in order to avoid biases.

## 3. Results

### 3.1. Selection of the size of **z** (D) and reproducibility of **A**

Our model was overfitted to the training data, and the inferred **A** was unstable with needlessly large *D*. Additionally, the model cannot reconstruct expression dynamics with insufficiently small values of *D*. Therefore, the selection of appropriate values for *D* is necessary, and we applied SCODE to training data and evaluated the validity of the optimized model on the basis of the RSS of independent test data for various values of *D* (*D* = 2, 4, 6, and 8). For each *D*, we executed SCODE 100 times independently, and the first, second, and third quantiles of the RSS values of test data are shown in Fig. 2(a). For every dataset, the median of RSS is almost saturated at *D* = 4.

**Figure 2:**
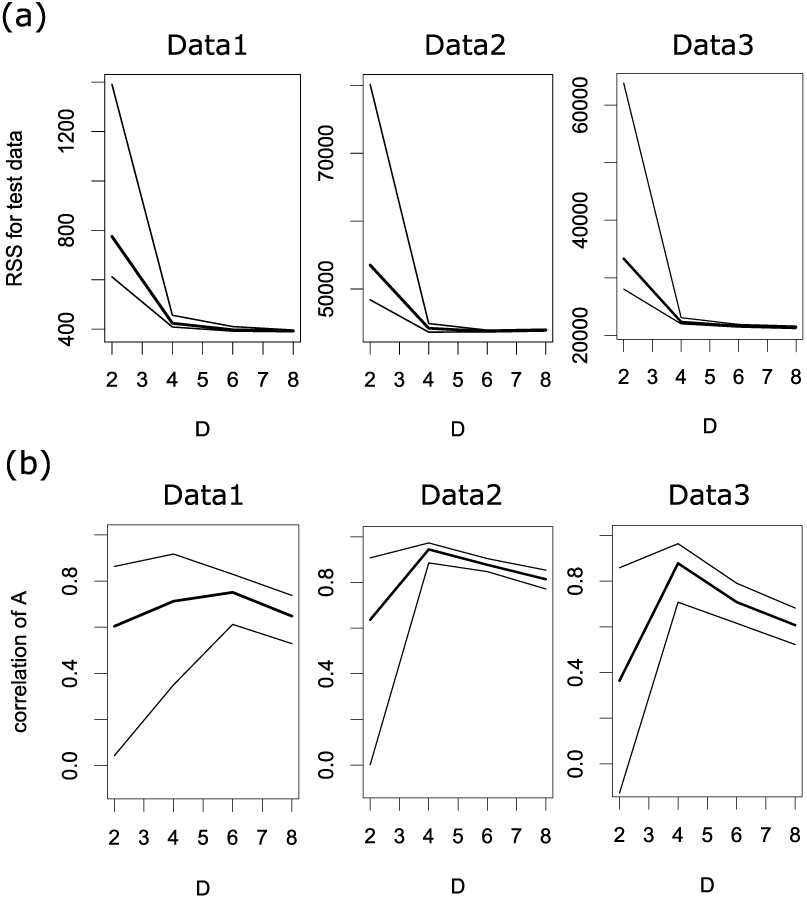
(a) The first, second, and third quantiles of the RSS values of test data (a) and the correlations among optimized **A** of the top 50 replicates (b) for each *D* (*D* = 2, 4, 6, and 8) for each dataset.

Because we used random sampling during optimization, we validated the reproducibility of the optimized **A**. We calculated the correlation coefficient among optimized **A** for the top 50 replicates (in ascending order of RSS values) of test data for each *D*. The corresponding first, second, and third quantiles of correlation coefficients are shown in Fig. 2(b). For *D* = 4, the medians of the correlation coefficients are 0.71, 0.94, and 0.88 for each dataset. The medians tend to decrease for large *D* because the matrix **A** is unstable with needlessly large *D*. The medians also decrease for small *D*, possibly because the optimized **A** is trapped in local optima. In summary, the correlations among replicates are high, and therefore, an optimized matrix **A** is stable for *D* = 4.

Because the RSS values for test data are almost saturated and the estimated **A** are stable with *D* = 4, we used *D* = 4 unless otherwise specified. For optimized **A** of each *D*, we used the mean of optimized **A** of the top 50 replicates, hereafter.

### 3.2. Validation of **A** optimization with simulation data

Next, we investigated whether SCODE can infer genuine **A** by using simulated data. Because the dynamics of **x** become unrealistic with randomly determined **A**, we used previously inferred **A** (for *D* = 4) as genuine **A** and simulated data with the same condition for each dataset (such as the same pseudo-time). We also added uniform random numbers (*ϵ* 𝜖 [−0.1, 0.1]) to simulated data as a noise term. We optimized **A** for each simulated dataset 100 times, and Fig. 3 shows the first, second, and third quantiles of the correlation coefficients between the genuine **A** and optimized **A** for each *D*. The medians are 0.70, 0.71, and 0.91 for *D* = 4, and 0.61, 0.48, and 0.49 for *D* = 6. Therefore, SCODE can accurately infer the genuine **A** with appropriate *D*, and can roughly infer **A** with slightly different *D* values unless we set extremely large or small *D*.

**Figure 3:**
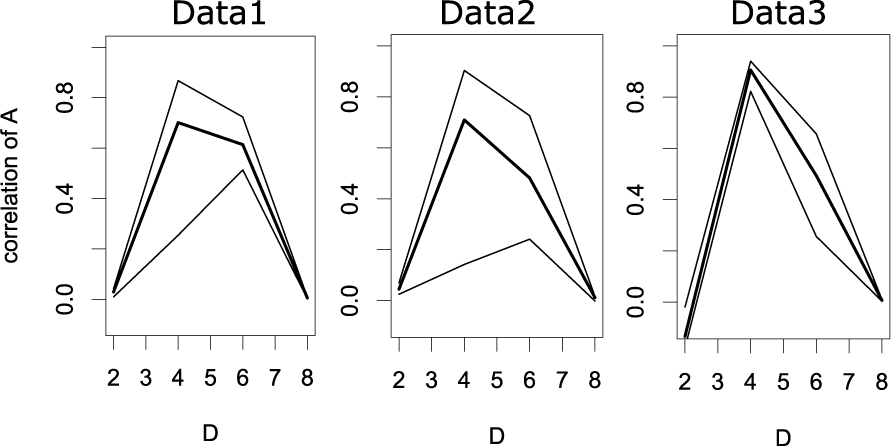
The first, second, and third quantiles of the correlation coefficients between genuine **A** and inferred **A** for each *D*.

### 3.3. Reconstruction of expression dynamics

Although RSS values for test data were almost saturated at *D* = 4, this does not necessarily mean that SCODE can successfully learn the dynamics. Next, we investigated whether the optimized ODE can accurately reconstruct observed expression dynamics to verify the optimization of SCODE (Fig 1(e)). For each set of dynamics, the initial values (**x** at *t* = 0) were set to the mean expression of 0-h or day 0 cells. At first, we compared the reconstructed dynamics with observed data in the principal component analysis (PCA) space (Fig 4). For every dataset, SCODE was able to reconstruct the dynamics with *D* ≥ 4.

**Figure 4:**
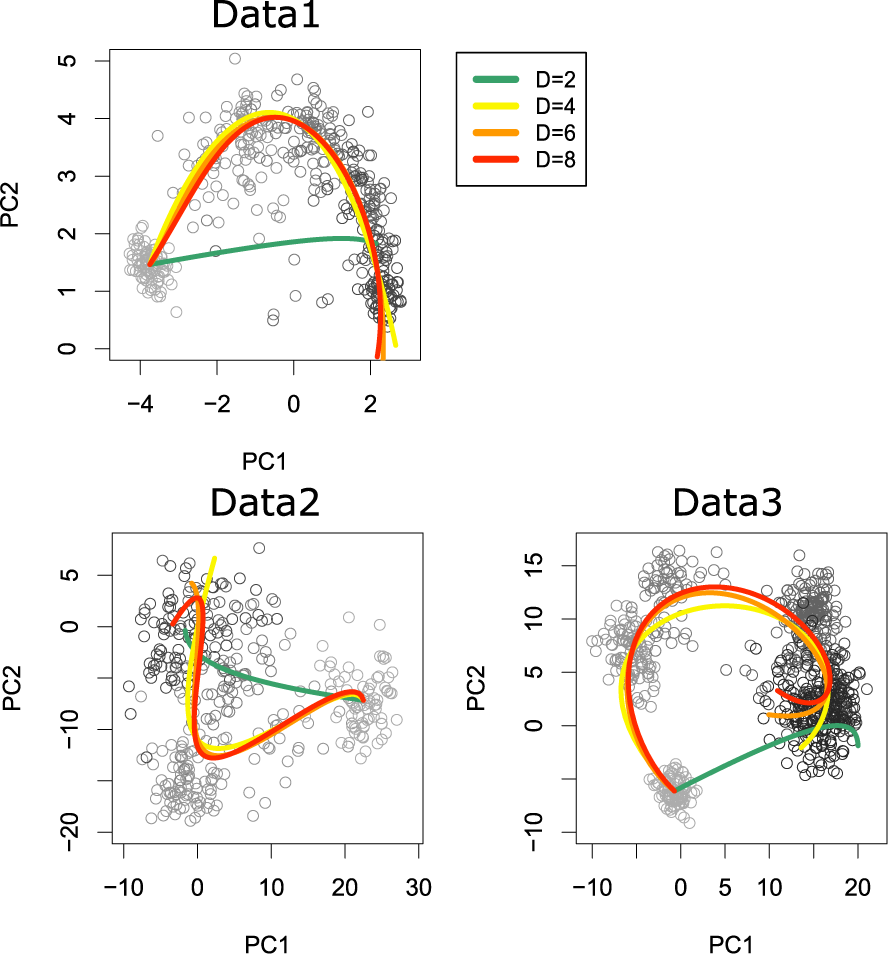
PCA of scRNA-Seq data for each dataset. Each circle represents a cell, and its color represents experimental time (from light gray to black). The reconstructed expression dynamics are projected onto PCA space and are represented by colored lines (green, yellow, orange, and red correspond to *D*=2, 4, 6, and 8, respectively).

Next, we compared the reconstructed dynamics with observed expression dynamics for some TFs (*Sox2*, *Utf1*, *Epas1*, and *Foxq1*) in Data1 (Fig 5). The analysis for every TF and dataset is described in the supplementary text. Although the reconstructed dynamics of SCODE with *D* = 2 differ from the observed data, the model with *D* ≥ 4 successfully reconstructed complicated dynamics, such as transient patterns. Therefore, we concluded that SCODE can successfully optimize **A** and learn the ODE of **x**

**Figure 5:**
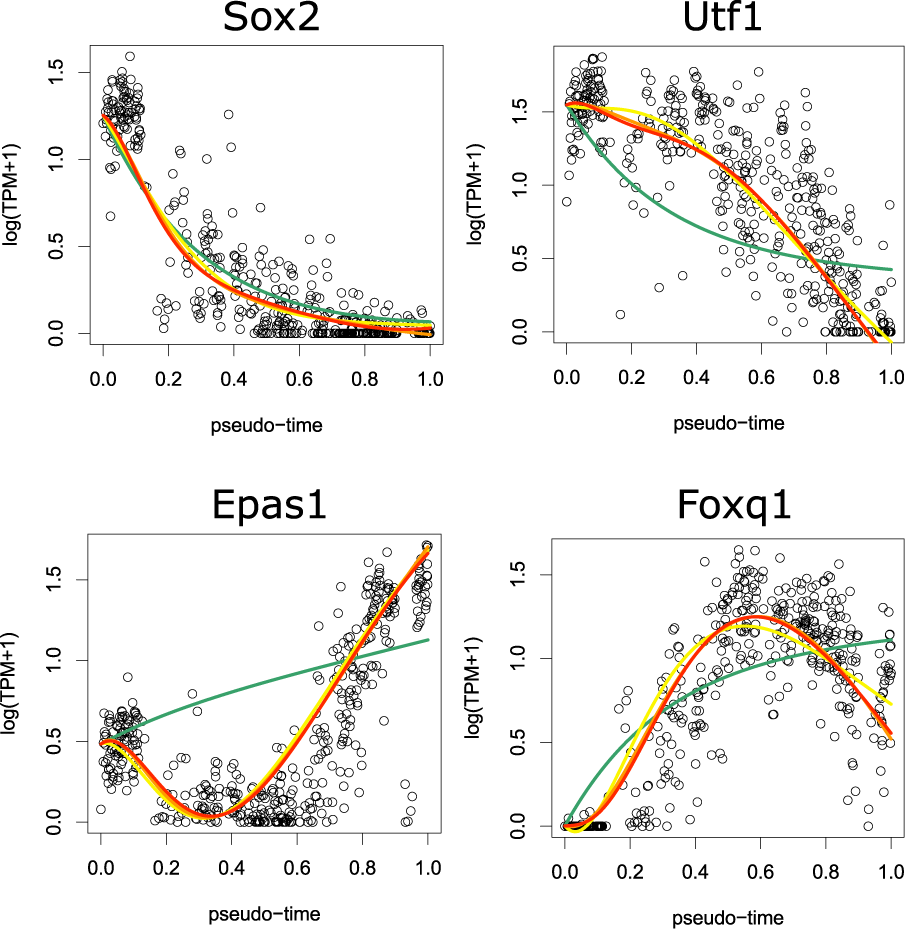
Observed expression of four TFs and reconstructed dynamics for each *D* (green, yellow, orange, and red correspond to *D*=2, 4, 6, and 8, respectively). The *x*-axis represents pseudo-time and *y*-axis represents log(TPM+1).

### 3.4. Validation of inferred network

We also evaluated the inferred network of each algorithm including the correlation network by comparing them to TF regulatory networks based on DNaseI footprints and TF-binding motifs (see section 2.4). Because the runtimes of Jump3 are significantly large for large numbers of cells, we used 25 cells at even intervals in the pseudo-time order as the data for Jump3. The AUC values of each method for each dataset are shown in Table 1.

**Table 1:** The AUC values of each method for each dataset. Cor is the correlation network.

|  | SCODE | lm | msgps | Cor | GENIE3 | Jump3 |
| --- | --- | --- | --- | --- | --- | --- |
| Data1 | 0.536 | 0.480 | 0.510 | 0.505 | 0.474 | 0.504 |
| Data2 | 0.581 | 0.489 | 0.516 | 0.492 | 0.472 | 0.492 |
| Data3 | 0.523 | 0.480 | 0.499 | 0.524 | 0.522 | 0.501 |

For Data1 and Data2, the AUC values of SCODE are significantly larger than those of the other algorithms. This is because our model considers the dynamics of expression and fully uses time information. Although Jump3 is also designed for time-course expression data, the AUC values are not high. This is because Jump3 is not designed for scRNA-Seq conducted during differentiation, but is designed for multiple time-course data. This suggests the necessity of a novel computational algorithm designed for scRNA-Seq data.

The performance of SCODE is second, but almost equal to the best performance for Data3. Given that the reconstructed path in PCA space is a little out of alignment for Data3 (Fig 4), our model based on linear ODEs might be slightly insufficient to describe the expression dynamics of Data3.

In summary, our algorithm can infer TF regulatory networks with high performance in comparison to other network inference algorithms, especially for Data1 and Data2. This results implies the importance of time parameters in network inference and the necessity of a novel network inference algorithm designed for scRNA-Seq data obtained during differentiation.

### 3.5. Runtimes

We investigated the runtime of each method and the runtimes for Data1 are shown in Table 2. The runtime of Jump3 is calculated using the data from 25 cells as stated above. The runtime of SCODE is 11 seconds and is significantly smaller than that of Jump3. Moreover, the runtime of SCODE is smaller than those of msgps and GENIE3, which do not consider time dynamics. These results show that SCODE can infer regulatory networks efficiently, even though it considers a time parameter in its model.

**Table 2:**
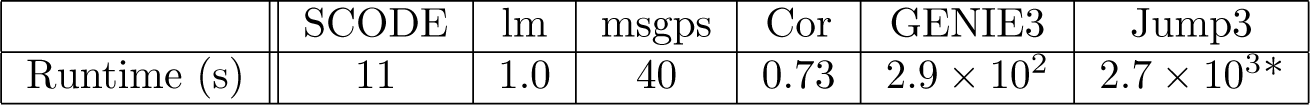
The runtimes of each method for Data1 (456 cells). The runtime of Jump3 is calculated using data from 25 cells. The computations were performed on a MacBook Pro equipped with a 3.1 GHz Intel Core i7 processor and 16 GB of 1867 MHz DDR3 RAM.

### 3.6. Network analysis

Lastly, we investigated the structure of the inferred regulatory network of Data1. At first, we defined the threshold *α* as the value of the 1000th largest absolute value in **A**, and we counted the number of positive edges (**A***_ij_* ≥ *α*) and negative edges (**A***_ij_* ≤ −*α*) for TF *j*. Figure 6(a) shows the total counts for each TF in decreasing order. About 39% of edges are included in the top 10 TFs, and this result implies the existence of key regulators for differentiation. Interestingly, most TFs mainly have either positive or negative edges, and this result suggests that TFs might mainly work as either activators or inhibitors in differentiation. This tendency was shared with Data3, but was not seen in Data2 (see supplementary text). This result might reflect a difference in the systems; Data1 and Data3 represent differentiation from ES cells, while Data2 represents direct reprogramming from MEF cells.

**Figure 6:**
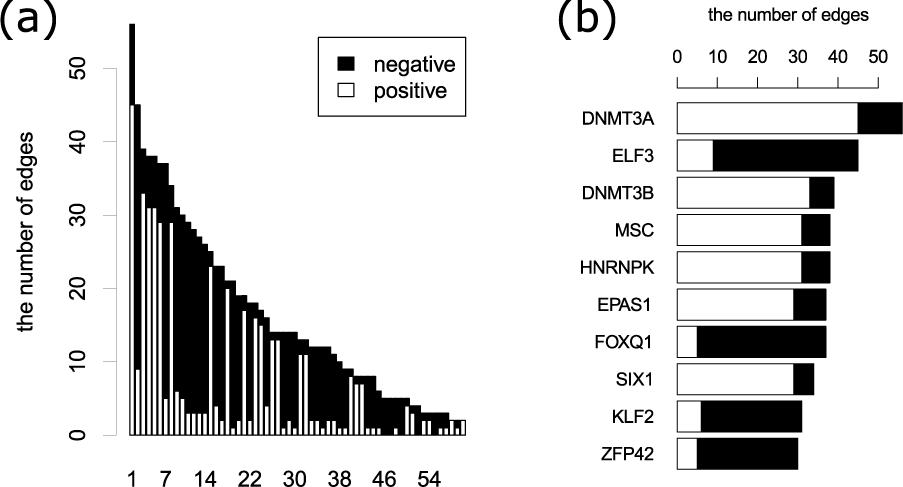
(a) Bar graph of positive and negative edges of each TF in decreasing order. For visibility, only the top 60 TFs are shown (see supplementary text for plot of all TFs). (b) Bar graph of the top 10 TFs.

We also visualized the top 10 TF results Fig 6(b). Interestingly, *Dnmt3a* and *Dnmt3b*, which are the de novo DNA methyltransferase have several positive edges. Data1 is derived from scRNA-Seq obtained from cells differentiating from mouse ES cells into PrE cells. To maintain the pluripotency of ES cells, *Dnmt3a* and *Dnmt3b* seem dispensable, and these genes must be unimportant for ES cells [36]. However, several studies have suggested the importance of *Dnmt3a* and *Dnmt3b* in differentiation. For example, these TFs restrict the lineage-specific function of TFs during differentiation via DNA methylation [37]. In addition, *Dnmt3a* is essential for hematopoietic stem cell differentiation and it seems to enhance differentiation by epigenetic silencing of multipotency genes [38]. Thus, *Dnmt3a* and *Dnmt3b* are known to affect differentiation based on de novo DNA methylation.

In this study, these genes were inferred to regulate several TFs positively. Because DNA methylation essentially silences expression, these TFs might be regulated positively indirectly via the inactivation of negative regulators of these TFs. Although the direct targets of *Dnmt3a* and *Dnmt3b* are obscure, our result suggests that they are the key regulators of this differentiation.

## 4. Discussion

The advancement of scRNA-Seq and the analysis of differentiation reconstruction and pseudo-time have elucidated differentiation mechanisms. The inference of regulatory networks associated with differentiation is necessary to further our understanding of differentiation and development. In the inference of regulatory networks, it is important to fully use pseudo-time information and expression dynamics. However, there are no efficient algorithms for inferring the regulatory networks of many TFs from continuous time expression data. Thus, we developed SCODE, an efficient algorithm based on linear ODEs. SCODE is based on the transformation of linear ODEs and linear regression, and the time complexity is significantly small.

We applied SCODE to three scRNA-Seq datasets during differentiation and showed that SCODE can successfully optimize ODEs so that these ODEs can reconstruct observed expression dynamics. In the validation of the inferred network, the AUC values of SCODE were higher than those of other methods in almost of all cases. The runtime of SCODE is significantly smaller than that of Jump3, which also infers networks from time-course data. Additionally, SCODE is faster than GENIE3, which does not use time information. These performance results show the efficiency of SCODE.

Single-cell sequencing technologies are developing rapidly, and the number of scRNA-Seq datasets produced from differentiating cells will therefore increase. Our novel and efficient method for inferring regulatory networks demonstrated high performance and will therefore enhance the analysis of regulatory networks.

Moreover, our model can reconstruct expression dynamics accurately. This means that we can simulate expression dynamics (such as those associated with the knockout of a TF) by using an optimized model, and such simulation-based analyses will be useful for many types of research, such as detection of drivers of differentiation. Thus, SCODE is useful not only for regulatory network inference, but also for various analyses using simulation, and therefore, our research is a promising computational tool for further single-cell sequence analyses.

## Acknowledgements

The authors thank Yohei Sasagawa, Hiroki Danno, Masashi Ebisawa, Mana Umeda, and Haruka Ozaki for assistance in this study. We also thank Tsukasa Fukunaga for critically reading the manuscript and Suguru Yaginuma for helpful discussions about our algorithm.

## Funding

This work was supported by CREST from Japan Science and Technology (JST), a Grant-in-Aid for JSPS Fellows, and JSPS KAKENHI Grant Number 16J05079.

